# Early AMPA receptor potentiation modifies synaptic maturation and disease progression in Rett models

**DOI:** 10.64898/2026.08.04.742773

**Authors:** Giuseppina De Rocco, Andrea de Donato, Marzia T Indrigo, Virginia Varotto, Martina Geusa, Stefano Taverna, Ingrid Cifola, Eva M Pinatel, Angelisa Frasca, Nicoletta Landsberger

## Abstract

Rett syndrome (RTT) is a severe neurodevelopmental disorder caused by mutations in *MECP2* and characterized by impaired neuronal maturation and synaptic dysfunction. Positive allosteric modulators of AMPA receptors (AMPAR-PAMs) have shown therapeutic promise in RTT models, but the determinants of treatment responsiveness remain unclear. Here, we evaluated the clinically advanced AMPAR-PAM CX1632 in *Mecp2*-null male and *Mecp2*-heterozygous female mice across developmental stages and treatment regimens. Therapeutic efficacy was strongly influenced by developmental stage, disease severity, and treatment schedule. Brief neonatal treatment produced long-lasting improvements in survival, disease progression, motor function, and cognition, whereas later intervention was markedly less effective in symptomatic null mice but remained beneficial in less severely affected heterozygous females. Repeated intermittent administration further enhanced selected benefits. Mechanistically, early CX1632 treatment induced sustained activation of neuronal and synaptic gene programs, restored synaptic organization and neuronal activity, and rescued AMPA receptor-mediated transmission weeks after drug withdrawal. These findings identify disease stage as a key determinant of responsiveness to AMPA receptor potentiation and support developmentally informed therapeutic strategies for *MECP2*-related disorders.

## Introduction

Mutations in the X-linked *MECP2* gene cause a broad spectrum of neurodevelopmental disorders, among which Rett syndrome (RTT) is the most prevalent and primarily affects females (Gold et al, 2024). RTT is characterized by an apparently normal early developmental period, typically lasting 6-18 months, followed by neurodevelopmental regression with loss of acquired motor, cognitive and communicative skills, and severe neurological symptoms, including intellectual disability, epilepsy, apraxia, progressive motor impairment and respiratory dysfunction (Gold et al, 2024, Fu et al. 2020).

*MECP2* encodes an epigenetic reader of methylated DNA with a central role in transcriptional regulation (Gold et al, 2024, Palmieri et al, 2023). Although broadly expressed, MeCP2 is highly abundant in neurons, where it is required for neuronal maturation and function (Skene et al, 2010, Kishi, Macklis, 2004). *Mecp2* loss-of-function mouse models, including the widely used *Mecp2*^tm1.1Bird^ null line (Guy et al, 2001), have shown that synapses and dendritic spines are particularly vulnerable to Mecp2 deficiency (Molina Calistro et al, 2025). Reduced spine density, altered spine morphology, and impaired neuronal arborization are hallmarks of RTT models, whereas neuronal number is largely preserved (Chapleau et al, 2009). In line with these synaptic abnormalities, *Mecp2-*deficient mouse models show alterations in synaptic transmission that affect circuit-level plasticity (Gold et al, 2024). Furthermore, symptomatic RTT mouse brains display a generalized reduction in neuronal activity, associated with altered excitatory/inhibitory balance, although region-specific exceptions exist, such as increased hippocampal activity (Molina Calistro et al, 2025, Kron et al, 2012, Calfa et al., 2015). Moreover, in several brain areas, long-term potentiation (LTP) and long-term depression (LTD)—key mechanisms of activity-dependent synaptic plasticity and learning and memory—are consistently impaired by *Mecp2* deficiency, together with neuronal responses to stimuli (Della Sala, Pizzorusso, 2014). Importantly, increasing evidence indicates that subtle anomalies are already present before regression. Indeed, from birth onwards, RTT patients often follow altered developmental trajectories and show delayed developmental milestones (Zhang et al, 2023). Accordingly, developing RTT neurons display defective maturation, impaired glutamatergic signaling and reduced neuronal activity (Xu et al, 2014, Bedogni et al, 2016, Cobolli Gigli et al, 2018). Because impaired glutamatergic signalling and reduced neuronal activity emerge early in RTT models, AMPA receptor (AMPAR) modulation may help enhance synaptic function during disease-relevant developmental windows. AMPAR positive allosteric modulators (PAMs), or ampakines, potentiate excitatory transmission without directly activating the receptor, thereby preserving the natural spatial and temporal signalling patterns of AMPARs (Radin et al, 2025, Kadriu et al, 2021). Interestingly, ampakines enhanced key features of synaptic function, including spine maturation and LTP, in various experimental systems (Baudry et al, 2012). Furthermore, previous studies using the ampakine CX546 have reported beneficial effects in RTT models (Ogier et al, 2007, Degano et al, 2014), supporting AMPAR positive modulation as a therapeutic strategy. Consistently, we showed that brief neonatal treatment with CX546 produces long-lasting benefits in *Mecp2*-null mice, suggesting that early enhancement of excitatory transmission can redirect altered neurodevelopmental trajectories (Scaramuzza et al, 2021).

However, CX546 has limited translational potential, and whether more clinically viable AMPAR modulators can produce sustained benefits in RTT models remains unknown. CX1632 is a potent AMPAR-PAM with preferential activity on GluA1-containing AMPA receptors and improved drug-like properties (Bretin et al, 2017, Bellingacci et al, 2025, Wilkinson et al, 2019). Yet, its molecular activity in cortical neurons, its efficacy in *Mecp2*-deficient mice, and the treatment windows required for durable therapeutic effects have not been defined.

We investigated whether CX1632 can engage AMPAR-associated signalling and rescue RTT-related phenotypes in *Mecp2*-null males and *Mecp2*-heterozygous females. We compared brief neonatal, late juvenile, and intermittent treatment regimens, and combined behavioural analyses with transcriptomic, synaptic, calcium imaging, and electrophysiological approaches.

Our findings indicate that CX1632 efficacy is strongly influenced by treatment timing, with early intervention producing the most sustained benefits during a developmental window in which RTT-related neuronal defects remain amenable to modulation.

## Results

### CX1632 activates GluA1/Akt signaling in cortical neurons and reveals preserved AMPAR availability in *Mecp2*-null mice

CX1632 is a potent AMPAR-PAM, whose activity depends on ligand binding to AMPA receptor (Bretin et al, 2017). The compound showed no toxicity in rat primary neurons and protected against glutamate-induced toxicity at 10 µM (Bretin et al, 2017). However, its molecular activity remains poorly defined. Since CX1632 preferentially modulates GluA1-containing AMPA receptors (Bretin et al, 2017), we first tested its effects in cultured mouse cortical neurons.

Based on the short half-life of the compound, wild type (WT) primary cortical neurons at DIV14 were treated with CX1632 (5 μM) for 1 hour and GluA1 phosphorylation was analysed by Western blot (WB; Fig. 1A). CX1632 significantly increased GluA1 phosphorylation at Ser845, without altering total GluA1 levels consistent with activation of GluA1-associated signalling (Fig. 1B). The treatment also increased Akt phosphorylation at Ser473, in line with previous evidence linking ampakines to Akt activation (Clarkson et al, 2015) (Fig. 1C).

**Figure 1:**
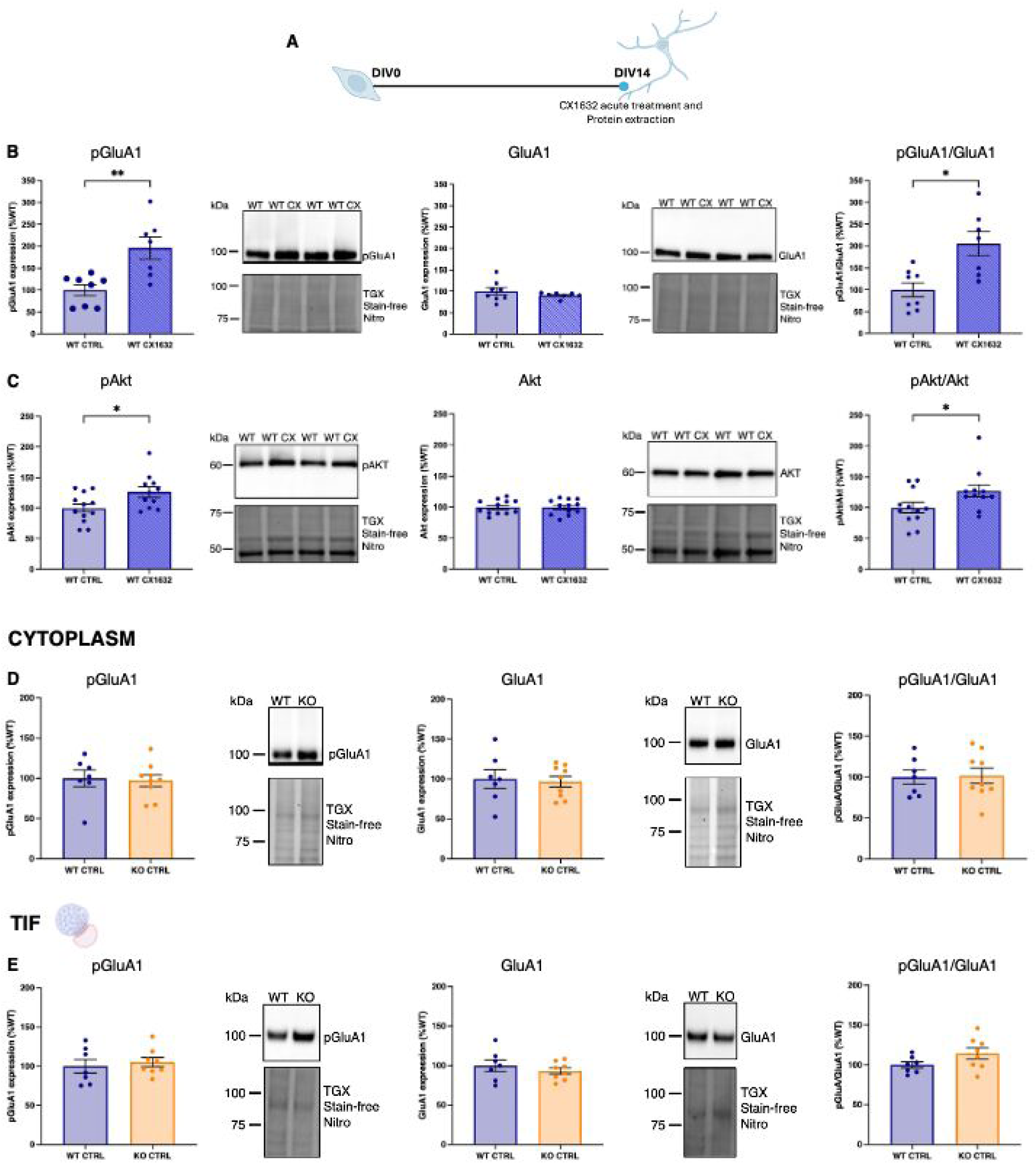
CX1632 activates GluA1/Akt signaling in cortical neurons, while GluA1 levels and phosphorylation are preserved in *Mecp2*-null mice. **(A)** Schematic representation of the *in vitro* CX1632 treatment protocol. **(B, C)** Quantification of pGluA1, total GluA1 and the pGluA1/GluA1 ratio (B) and pAkt, total Akt and pAkt/Akt ration (C) in untreated and CX1632-treated WT cortical neurons. Representative Western blots are shown on the right. Total protein content, visualized by TGX Stain-Free technology (Bio-Rad), was used for normalization. Each dot represents one embryo from three independent neuronal preparations. Statistical significance was assessed using the Mann-Whitney test and t-test (*: p-value<0.05, ***: p-value<0.001, ****: p-value<0.0001). Data are presented as mean ± SEM. **(D, E)** Quantification of pGluA1, total GluA1 and the pGluA1/GluA1 ratio in the cytosolic fraction and Triton-insoluble fraction (TIF), respectively, from P40 WT and *Mecp2*-null mice. Representative Western blots of pGluA1 and total GluA1 are shown on the right. Total protein content, visualized by TGX Stain-Free technology (Bio-Rad), was used for normalization. Each dot represents one animal. Sample sizes: WT=7, KO=9. Statistical significance was assessed using Mann-Whitney test test. Data are presented as mean ± SEM.

We then analysed total GluA1 and pSer845-GluA1 levels in cerebral cortices from adult (P40) WT and *Mecp2*-null (KO) male mice. Both the cytoplasmic fraction and the Triton-Insoluble Fraction (TIF), used as a synapse-enriched proxy for postsynaptic protein complexes (Gardoni et al, 1998), were examined. No genotype-dependent differences were detected in either fraction, indicating comparable basal GluA1 expression and phosphorylation in WT and symptomatic KO cortices (Fig. 1D,E).

Together, these data showed that CX1632 activates GluA1-associated signalling in cortical neurons and that *Mecp2*-null cortices retain molecular substrates compatible with AMPAR-PAM responsiveness.

### Early CX1632 administration produces long-lasting therapeutic benefits in *Mecp2-*mutant mice

We previously showed that brief neonatal treatment with the non-translational ampakine CX546 produces long-lasting benefits in *Mecp2-*null mice (Scaramuzza et al, 2021). We therefore tested the more clinically viable ampakine CX1632, using the same regimen (Scaramuzza et al, 2021). WT and KO male mice were injected daily with CX1632 (3 mg/kg) or vehicle from P3 to P9, and well-being and behavior were assessed by observers blinded to treatment group (Fig. 2A).

**Figure 2:**
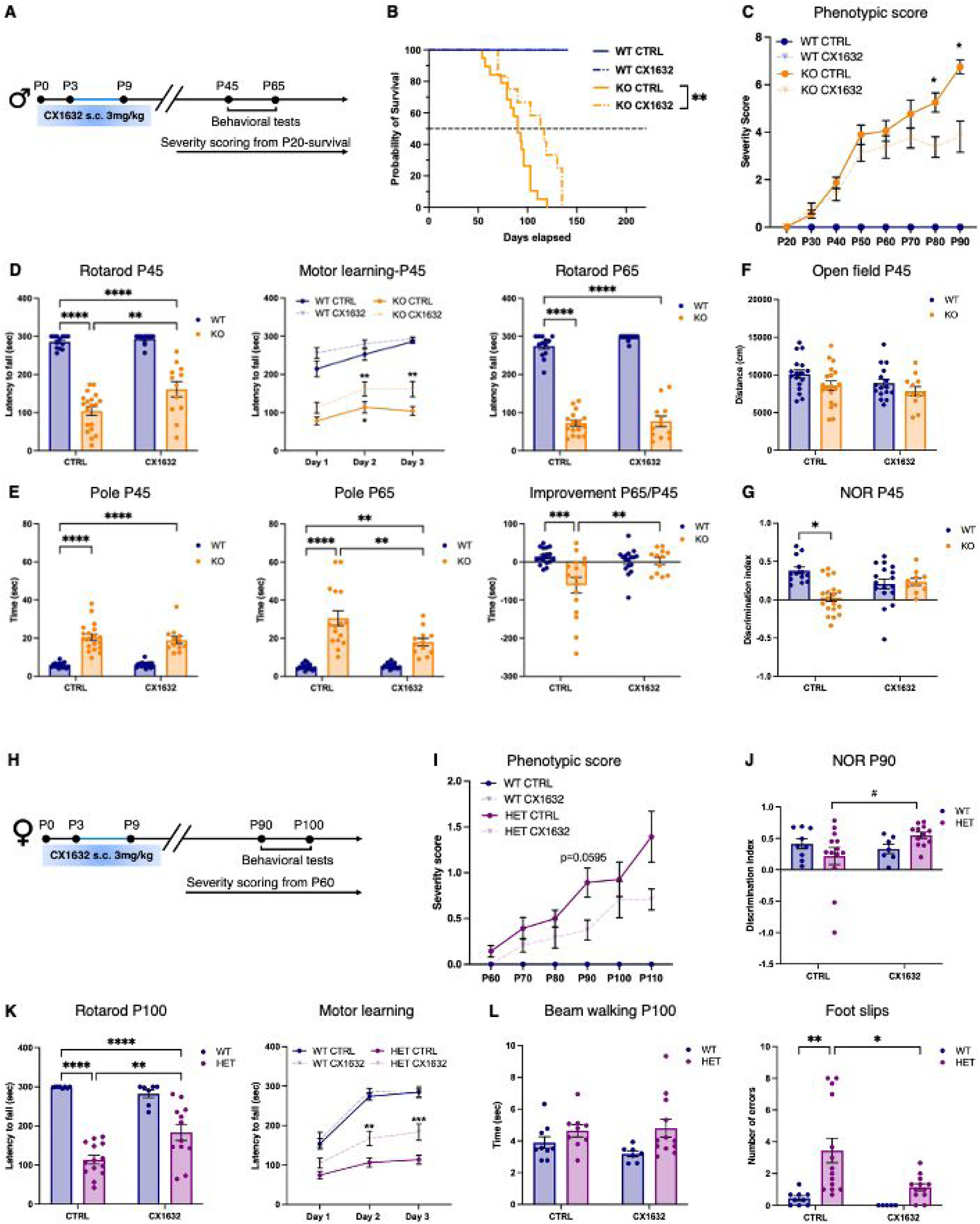
Early CX1632 treatment ameliorates behavioural deficits in *Mecp2*-null and HET mice. **(A)** Schematic representation of the early CX1632 treatment protocol. Male mice received daily injection of CX1632 or vehicle from P3 to P9. Severity score was assessed twice weekly from P20, and behavioural tests were performed at P45 and P65. **(B)** Kaplan–Meier survival curves showing prolonged survival in CX1632-treated *Mecp2*-null mice compared with vehicle-treated KO controls. **(C)** Longitudinal phenotypic severity scores from P20 to P90, based on general condition, mobility, hindlimb clasping, tremor and gait. (**D**) Rotarod performance at P45 and P65, expressed as latency to fall during the final trial, together with the learning curve across three days at P45. **(E)** Pole-test performance at P45 and P65, expressed as the time required to descend the pole. The longitudinal analysis shows the change in performance of individual mice between P45 and P65. **(F)** Distance travelled during the open-field test at P45. **(G)** Discrimination index in the novel object recognition test at P45. Each dot represents one animal. Sample sizes before outlier exclusion were: WT, n = 17; CX1632-treated WT, n = 17; KO, n = 19; and CX1632-treated KO, n = 12. Outliers were excluded using the ROUT test as described in the Methods. Statistical significance was assessed by two-way ANOVA followed by Tukey’s multiple-comparisons test, and by the log-rank (Mantel–Cox) test for survival analysis in (B).*: p-value<0.05, **: p-value<0.01, ***: p-value<0.001, ****: p-value<0.0001. Data are presented as mean ± SEM. **(H)** Schematic representation of the early CX1632 treatment protocol in female mice. WT and HET females received daily injections of CX1632 or vehicle from P3 to P9. Phenotypic severity was assessed weekly from P60, and behavioural tests were performed at P90 and P100. **(I)** Longitudinal phenotypic severity scores from P60 to P110, based on general condition, mobility, hindlimb clasping, tremor and gait. **(J)** Discrimination index in the novel object recognition test at P90. **(K)** Rotarod performance at P100, expressed as latency to fall during the final trial, together with the learning curve. **(L)** Beam-walking performance at P100, expressed as traversal time and number of foot slips. Each dot represents one animal. Sample sizes before outlier exclusion were: WT, n = 9; CX1632-treated WT, n = 7; HET, n = 14; and CX1632-treated HET, n = 12. Outliers were excluded using the ROUT test as described in the Methods. Statistical significance was assessed by two-way ANOVA followed by Tukey’s multiple-comparisons test. *P < 0.05, **P < 0.01, ***P < 0.001, ****P < 0.0001; #P = 0.0648. Data are presented as mean ± SEM.

CX1632 significantly increased survival of KO mice by approximately 30%, with benefits persisting for more than 100 days after treatment withdrawal (Fig. 2B). Phenotypic scoring (Guy et al, 2007) showed slower disease progression in CX1632 treated KO mice as compared to untreated controls, becoming evident around P55, 45 days after the last injection, and significant at P80 (Fig. 2C). Itemized scores indicated earlier effects on general condition and tremor, while mobility improved only at later stages (Fig. EV1A-E).

Motor performance was assessed by rotarod and pole test. At P45, CX1632-treated KO mice performed significantly better than vehicle-treated KO mice, although they did not reach WT levels (Fig. 2D). Analysis of the motor learning curve further indicated that treated KO mice continued to improve across trials, unlike untreated KO mice. This amelioration was no longer detectable at P65. Conversely, the pole test, which primarily assesses motor coordination and balance, showed improved performance in CX1632-treated KO mice only at P65 (Fig. 2E). Longitudinal analysis further suggested that early treatment delayed motor decline.

We evaluated cognitive function at P45 using the Novel Object Recognition (NOR) test. Open field analysis excluded major locomotor confounds (Fig. 2F). We observed that CX1632-treated KO mice did not show the short-term memory impairment occurring in untreated KO mice (Fig. 2G). Importantly, no detectable drug effects were observed in WT mice.

We next tested the same neonatal regimen in *Mecp2^+/-^* heterozygous (HET) females, which represent a more genetically relevant model of RTT. WT and HET mice were treated from P3 to P9 and analyzed from P60 onward (Fig. 2H). Since HET mice do not show reduced survival during the first year of life, survival was not assessed. Despite their milder and more heterogeneous phenotype, CX1632-treated HET mice showed slower disease progression than untreated HET controls (Fig.2I). In the NOR test, treated HET mice showed a trend toward improved short-term memory, although control HET mice were not significantly impaired compared with WT littermates (Fig. 2J). Locomotor activity was comparable across groups, excluding motor confounds at this age (Fig. EV1K).

Motor function was tested around P100, nearly 90 days after the last injection. CX1632-treated HET mice showed significantly improved rotarod performance compared with vehicle-treated HET mice and improved across trials (Fig.2K). In the beam-walking test, which assesses fine motor coordination and balance and is particularly sensitive to hindlimb function, CX1632 rescued the increased number of foot slips observed in control HET mice (Fig. 2L).

Overall, brief neonatal CX1632 treatment produced robust and long-lasting behavioural benefits in RTT mouse models, persisting for months after drug withdrawal and without detectable effects in WT littermates.

### Treatment timing and regimen shape CX1632 efficacy in juvenile *Mecp2-*null mice

Because RTT is usually diagnosed after regression, around 2-4 years of age (Ferreira et al, 2025), we next tested a later 7-day CX1632 treatment. *Mecp2-*null males were treated from P28 to P34, when overt mild symptoms first emerge in this model, corresponding approximately to a later juvenile stage in humans (Fig. 3A).

**Figure 3:**
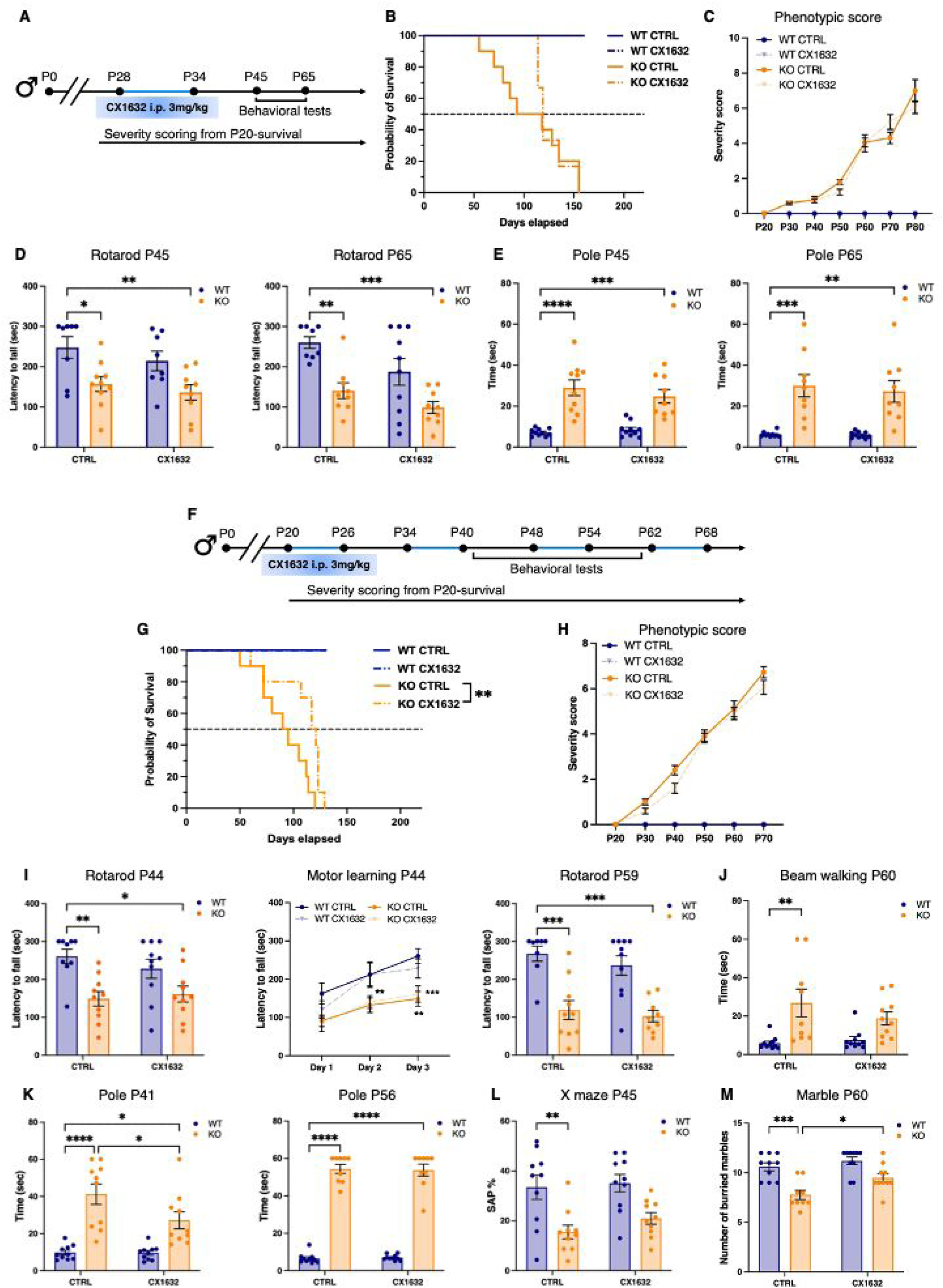
Late and intermittent juvenile CX1632 treatment produces regimen- and time-dependent effects in *Mecp2*-null male mice. **(A)** Schematic representation of the CX1632 treatment protocol. Mice received CX1632 or vehicle from P28 to P34 once daily. Phenotypic severity was assessed twice weekly from P20, and behavioral tests were performed between P45 and P65. **(B)** Kaplan–Meier survival curves showing no significant difference between CX1632-treated *Mecp2*-null mice and vehicle-treated KO controls. **(C)** Longitudinal phenotypic severity scores from P20 to P80, based on general condition, mobility, hindlimb clasping, tremor and gait. **(D)** Rotarod performance at P45 and P65, expressed as latency to fall during the final trial. **(E)** Pole-test performance at P45 and P65, expressed as total test completion time. Each dot represents one animal. Sample sizes before outlier exclusion were: WT, n = 8; CX1632-treated WT, n = 10; KO, n = 10; and CX1632-treated KO, n = 9. Statistical significance was assessed by two-way ANOVA followed by Tukey’s multiple-comparisons test and by the log-rank (Mantel–Cox) test for survival analysis in (B). *P < 0.05, **P < 0.01, ***P < 0.001, ****P < 0.0001. Data are presented as mean ± SEM. **(F)** Schematic representation of the intermittent CX1632 treatment protocol. Mice received CX1632 or vehicle every other week from P20 to P68. Phenotypic severity was assessed twice weekly from P20, and behavioural tests were performed between P41 and P45 and between P55 and P61. **(G)** Kaplan– Meier survival curves showing prolonged survival in CX1632-treated *Mecp2*-null mice compared with vehicle-treated KO controls. **(H)** Longitudinal phenotypic severity scores from P20 to P70, based on general condition, mobility, hindlimb clasping, tremor and gait. **(I)** Rotarod performance at P44 and P59, expressed as latency to fall during the final trial, together with the learning curve across three days at P44. **(J)** Beam-walking performance at P60, expressed as beam traversal time. **(K)** Pole-test performance at P41 and P56, expressed as total test completion time. **(L)** Percentage of spontaneous alternations in the X maze at P45. **(M)** Number of marbles buried after 30 min at P60. Each dot represents one animal. Sample sizes before outlier exclusion were: WT, n = 10; CX1632-treated WT, n = 10; KO, n = 10; and CX1632-treated KO, n = 10. Statistical significance was assessed by two-way ANOVA followed by Tukey’s multiple-comparisons test and by the log-rank (Mantel–Cox) test for survival analysis in (G). *P < 0.05, **P < 0.01, ***P < 0.001, ****P < 0.0001. Data are presented as mean ± SEM.

In contrast to early treatment, late CX1632 treatment did not significantly extend mean lifespan. However, vehicle-treated KO mice began to die at 55 days of age, with a mean survival of 105 days, whereas no CX1632-treated KO mice died before P114, after which mortality rapidly increased (Fig. 3B). Consistent with this delayed mortality onset, vitality differed between P80 and P120. Despite this trend, phenotypic scores progressed similarly in both KO groups (Fig. 3C; Fig. EV1G-K), and pole and rotarod tests showed no measurable treatment benefit (Fig. 3D,E).

This limited efficacy may reflect the lower ability of CX1632 to induce long-lasting functional remodelling in a later juvenile brain, where plasticity persists but is increasingly biased toward circuit refinement, stabilization and synaptic pruning, potentially constraining the reshaping of early neuronal wiring programs (Moyer, Zuo, 2018, Faust et al, 2021, Mallya et al, 2019). We thus tested CX1632 during an earlier juvenile phase characterized by heightened experience-dependent plasticity and ongoing refinement of excitatory and inhibitory circuits (Gordon, Stryker, 1996). To enhance efficacy while reducing continuous exposure, WT and KO mice received CX1632 every other week starting at P20. Behavioral tests were performed during off-treatment weeks, while severity scores and survival were monitored throughout (Fig. 3F).

This intermittent juvenile regimen significantly increased KO survival by approximately 30%, with mean lifespan rising from 92.5 days in vehicle-treated KO mice to 119 days in CX1632-treated KO mice (Fig. 3G). However, overall phenotypic scores were not improved (Fig. 3H; Fig. EV1L-P).

Behavioral benefits were modest but detectable. CX1632 did not improve rotarod performance (Fig.3I), but the beam-walking deficit observed in control KO mice was no longer evident in CX1632-treated KO mice (Fig. 3J). CX1632 also improved pole test performance at P41, but not at P56 (Fig.3K). In the X-maze spontaneous alternation test, CX1632 abolished the working-memory deficit seen in vehicle-treated KO mice (Fig. 3L). Finally, CX1632 fully rescued marble burying performance at P60, indicating improved environment-directed exploratory behavior in treated *Mecp2*-null mice (Frasca et al, 2024, Pozzer et al, 2025) (Fig. 3M).

Together, these experiments indicated that CX1632 can provide therapeutic benefits in young *Mecp2-* null male mice when administered through a prolonged intermittent regimen, whereas a brief later juvenile treatment has limited efficacy.

### Late intervention reveals preserved therapeutic responsiveness in juvenile HET females

The limited efficacy of P28-P34 CX1632 treatment in KO mice could reflect reduced brain plasticity or the cumulative burden of abnormalities established since embryonic development (Bedogni et al, 2016). To distinguish between these possibilities, we tested the same treatment window in HET females (Fig. 4A), which do not yet display overt phenotypic abnormalities (Cobolli Gigli et al, 2016). Although CX1632 did not improve the phenotypic score (Fig. 4B), behavioral testing revealed efficacy comparable to early P3–P9 treatment. CX1632 normalized the rotarod and beam-test deficits observed in vehicle-treated HET females relative to WT controls (Fig.4 C,D). As no genotype-dependent deficit emerged in the NOR test, mice were subsequently assessed in a novelty-exposure paradigm, in which an object was introduced into an empty arena after 10 minutes of habituation. During the first 5 min of object exposure, vehicle-treated HET females spent more time actively exploring the object than WT controls, despite a comparable number of exploratory bouts (Fig. 4E). CX1632 normalized exploration time without rescuing the locomotor deficit observed in the arena (Fig. 4E,F). Thus, the increased exploration time likely reflected prolonged object-directed exploration rather than more frequent approaches, possibly indicating altered habituation, perseveration, or reduced disengagement from a salient stimulus. Its normalization without locomotor rescue suggests a selective effect of CX1632 on the temporal organization of exploratory behavior.

**Figure 4:**
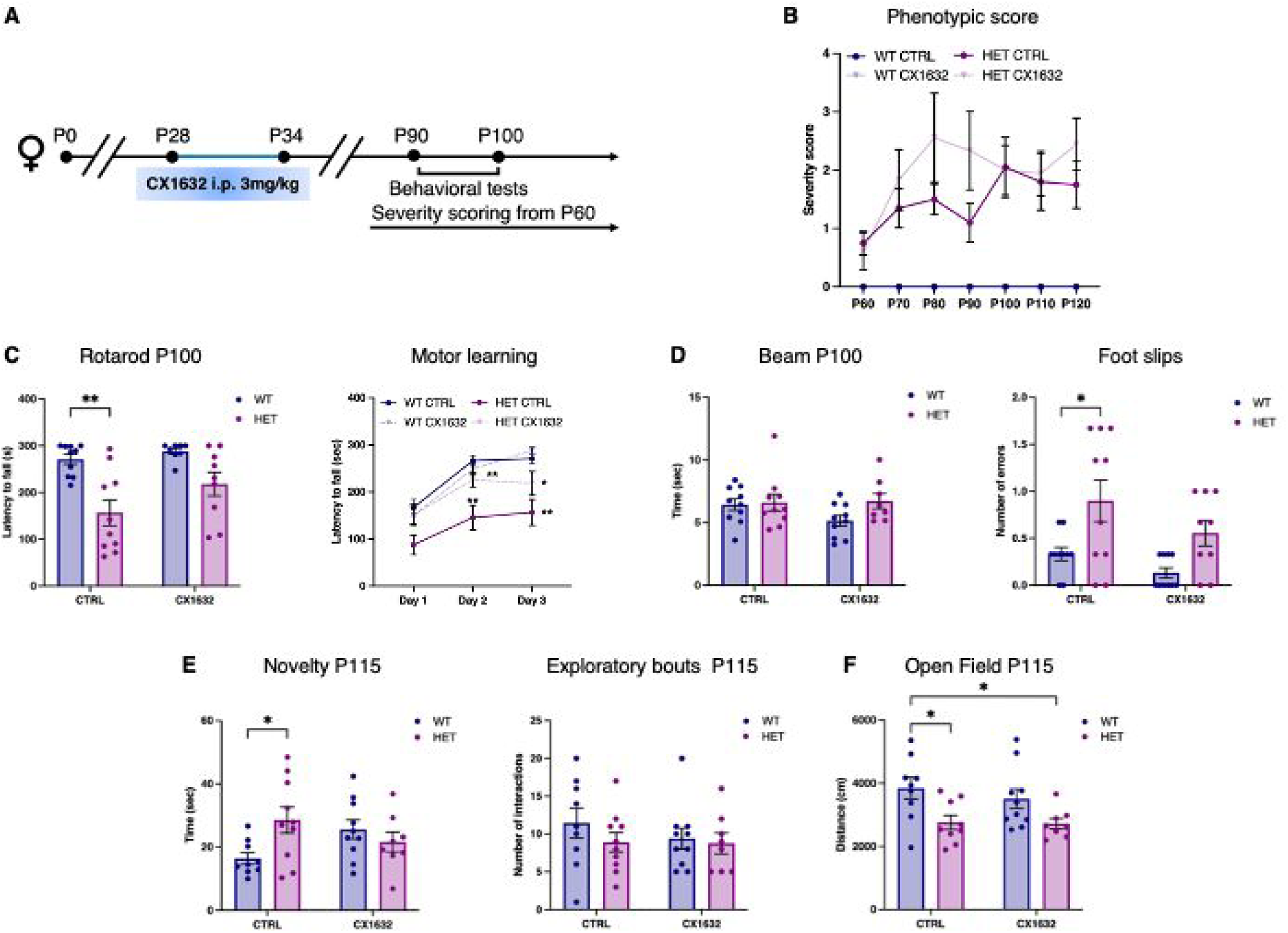
Juvenile HET females retain therapeutic responsiveness to late CX1632 treatment. **(A)** Schematic representation of the CX1632 treatment protocol. WT and HET females received daily intraperitoneal injections of CX1632 or vehicle from P28 to P34. Phenotypic severity was assessed from P60, and behavioral tests were performed in between P100 and P115. **(B)** Longitudinal phenotypic severity scores from P60 to P120, based on general condition, mobility, hindlimb clasping, tremor and gait. **(C)** Rotarod performance at P100, expressed as latency to fall during the final trial, together with the motor-learning curve across three consecutive testing days. **(D)** Beam-walking performance at P100, expressed as traversal time and number of foot slips. **(E)** Time spent actively exploring the object and number of discrete exploratory bouts during the first 5 min of object exposure at P115. **(F)** Distance travelled during the open-field habituation phase at P115. Each dot represents one animal. Sample sizes before outlier exclusion were: WT, n = 10; CX1632-treated WT, n = 10; HET, n = 10; and CX1632-treated HET, n = 9. Outliers were excluded using the ROUT test as described in the Methods. Statistical significance was assessed by two-way ANOVA followed by Tukey’s multiple-comparisons test. *P < 0.05, **P < 0.01. Data are presented as mean ± SEM.

### Early intermittent CX1632 treatment strengthens therapeutic efficacy in *Mecp2*-null mice

Since earlier CX1632 administration was associated with greater therapeutic efficacy, we next asked whether an intermittent regimen starting at P3 could further enhance its effects in KO mice. Based on the sustained efficacy observed after neonatal treatment, mice received repeated cycles consisting of seven days of CX1632 administration followed by two weeks off treatment (Fig. 5A).

**Figure 5:**
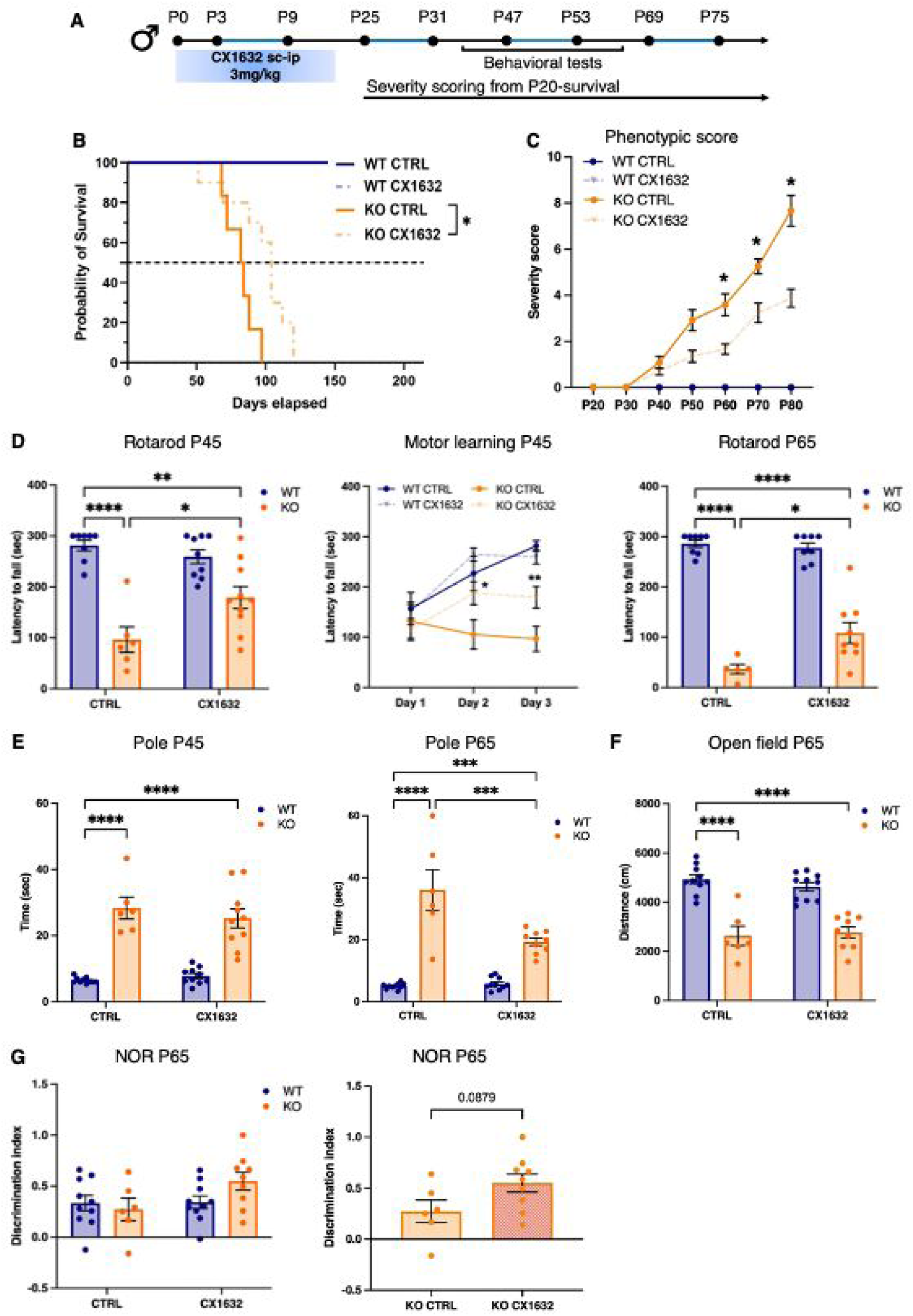
Early intermittent CX1632 treatment strengthens and prolongs behavioural benefits in *Mecp2*-null mice. **(A)** Schematic representation of the intermittent CX1632 treatment protocol. Mice received daily CX1632 or vehicle from P3 to P9, followed by repeated seven-day treatment cycles separated by two-week off-treatment intervals. Phenotypic severity was assessed twice weekly from P20, and behavioral tests were performed between P45 and P65. **(B)** Kaplan–Meier survival curves showing prolonged survival in CX1632-treated *Mecp2*-null mice compared with vehicle-treated KO controls. **(C)** Longitudinal phenotypic severity scores from P20 to P80, based on general condition, mobility, hindlimb clasping, tremor and gait. **(D)** Rotarod performance at P45 and P65, expressed as latency to fall during the final trial, together with the learning curve at P45. **(E)** Pole-test performance at P44 and P65, expressed as total test completion time. **(F)** Distance travelled during the open-field test at P65. (**G, H)** Discrimination index in the novel object recognition test at P65. Each dot represents one animal. Sample sizes before outlier exclusion were: WT, n = 10; CX1632-treated WT, n = 10; KO, n = 6; and CX1632-treated KO, n = 10. Statistical significance was assessed by two-way ANOVA followed by Tukey’s multiple-comparisons test and by the log-rank (Mantel–Cox) test for survival analysis in (**B**). *P < 0.05, **P < 0.01, ***P < 0.001, ****P < 0.0001. Data are presented as mean ± SEM.

This regimen prolonged survival of treated KO mice by approximately 30%, with a mean lifespan of 120 days (Fig. 5B). Phenotypic scoring revealed a consistent and significant improvement from P60 onward (Fig. 5C), an effect that emerged 20 days earlier than in mice treated only from P3 to P9. Analysis of itemized scores further confirmed a slower progression of most phenotypic features in intermittently treated KO mice (Fig. EV1, compare panels A-E and Q-U).

Consistently, CX1632-treated KO mice showed significant improved rotarod performance at both P45 and P65 (Fig. 5D), while the pole test revealed a significant benefit only at P65 (Fig. 5E). Short-term memory was assessed using the NOR test. Open field analysis indicated reduced locomotor activity in KO mice, consistent with a motor phenotype (Fig. 5F). In the NOR test, however, two-way ANOVA did not detect a significant baseline impairment between vehicle-treated WT and KO mice, indicating the absence of a robust recognition-memory deficit in this cohort (Fig. 5G). Nevertheless, a direct comparison of vehicle- and CX1632-treated KO mice showed a trend toward improved discrimination performance following treatment (p=0.089), suggesting a possible beneficial effect of CX1632 (Fig. 5G).

Collectively these data indicated that the efficacy of early CX1632 treatment can be further enhanced by repeated intermittent administration.

### Early CX1632 treatment induces persistent up-regulation of neuronal and synaptic pathways in KO prefrontal cortex

We next investigated the molecular mechanisms potentially underlying the sustained efficacy of brief early CX1632 treatment. WT and KO mice were treated with CX1632 or vehicle from P3 to P9, and bulk RNA sequencing was performed on the prefrontal cortex at P30, 21 days after the last administration. This delayed time point was selected to avoid acute drug-induced transcriptional responses and to focus on persistent molecular changes potentially associated with long-lasting behavioural benefits observed after treatment withdrawal.

The prefrontal cortex was selected because cortical dysfunction is central to RTT pathophysiology and has been extensively characterized in *Mecp2*-null mice (Frasca et al, 2020, Pozzer et al, 2025. Dani et al, 2005, Urdinguio et al, 2008). Moreover, analysis of a restricted cortical region was expected to reduce transcriptional heterogeneity compared with whole-cortex profiling, while preserving relevance to disease-associated cortical circuits.

Six animals per group were analysed. Sequencing yielded at least 30M read pairs/sample, with ≥92% uniquely mapped reads to the mouse genome and 25M read pairs assigned to protein-coding genes.

Principal component analysis (PCA) showed overall sample separation by treatment and genotype (Fig. EV2A). However, three CX1632-treated KO littermates clustered apart from the other treated KO mice and closer to vehicle-treated KO samples. Since this litter-specific clustering could reflect reduced treatment responsiveness, a faster loss of treatment-induced molecular effects after drug withdrawal, or other litter-dependent sources of variability, the entire litter was conservatively excluded from the analysis to avoid potential litter-specific confounding. The final dataset showed a clear segregation of samples according to genotype and treatment (Fig. 6A).

**Figure 6:**
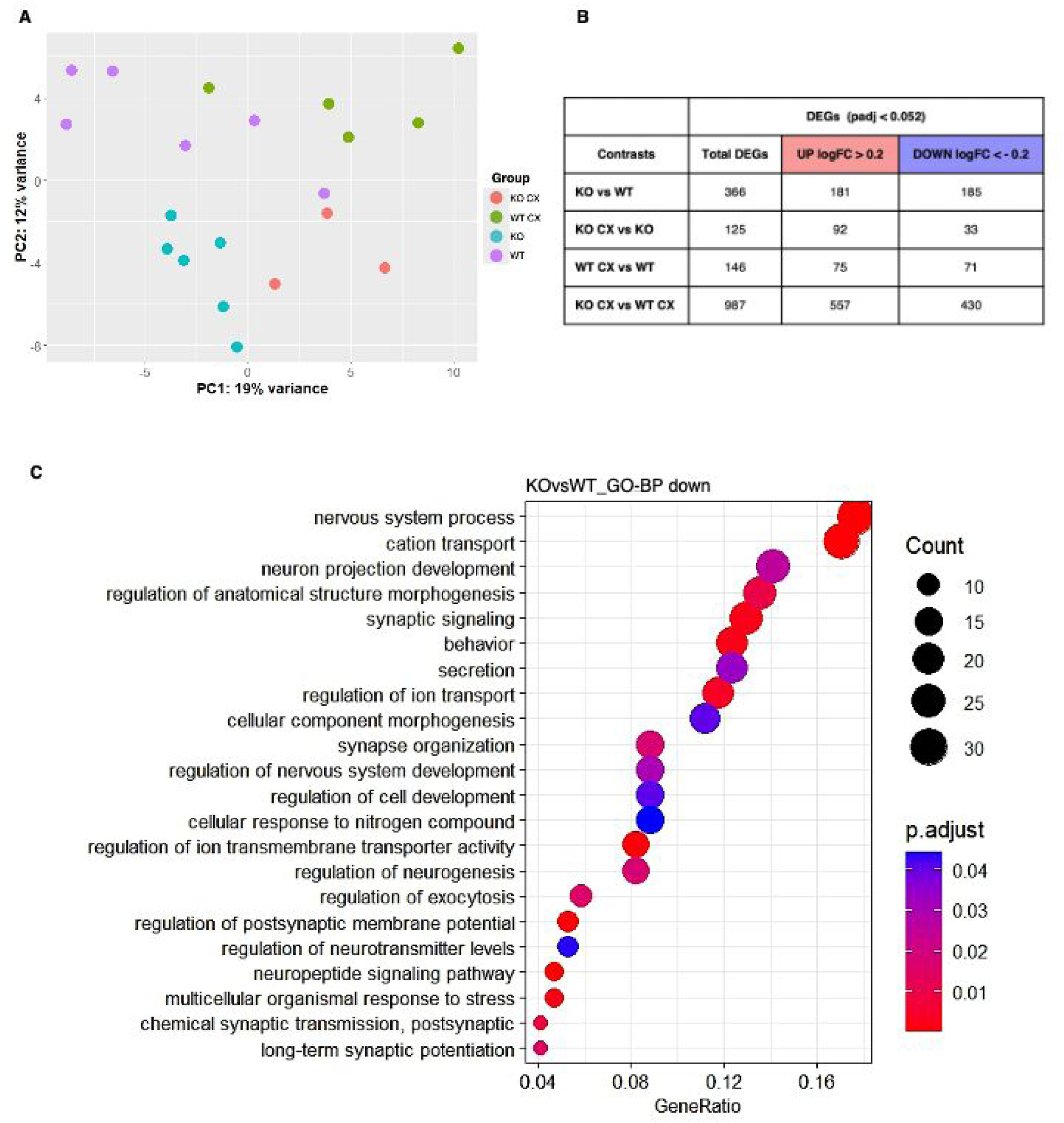
Early transcriptional alterations in *Mecp2*-null prefrontal cortex. **(A)** Principal component analysis of RNA-seq samples from the four experimental groups: vehicle-treated WT, vehicle-treated KO, CX1632-treated WT and CX1632-treated KO mice. **(B)** Number of differentially expressed genes for each comparison, defined by adjusted *P* < 0.052 and |log₂FC| > 0.2. **(C)** Functional enrichment analysis of DEGs identified in KO versus WT mice. The bubble plot shows selected Gene Ontology biological process terms enriched among downregulated genes. Bubble size indicates the number of DEGs contributing to each term, and color indicates enrichment significance.

At P30, when KO mice were only mildly symptomatic, KO vs. WT differential expression analysis identified 366 DEGs, indicating modest transcriptional dysregulation (Fig. 6B, and Supplementary File 1 for the complete annotated DEG lists). Interestingly, early CX1632 treatment altered a similar number of genes in KO and WT mice relative to their untreated controls, although changes in treated KO mice were predominantly up-regulated. However, only six treatment-responsive DEGs were shared between KO and WT mice, and DEGs in treated WT mice showed limited functional enrichment. Thus, CX1632 elicited a distinct and functionally more coherent transcriptional response in the *Mecp2*-deficient brain.

Functional enrichment analysis of DEGs using GO categories and pathway collections showed that in KO *vs.* WT mice most enriched processes were down-regulated and related to ion transport, neurogenesis, synaptic signalling, and synapse organization (Fig. 6C, and Supplementary File 2), as well as neuron projection, pre-synapse, ion channel complex, and somatodendritic compartment (Fig. EV2B, and Supplementary File 2). Reactome analyses revealed down-regulation of genes involved in GPCR signalling, nervous system development and axon guidance (Fig. EV2C and Supplementary File 2). Overall, these findings are consistent with known defects in neuronal development, maturation and synaptic functions in RTT models.

Interestingly, in treated vs. untreated KO mice, nearly all enriched processes were up-regulated and largely involved the same functional categories and cellular compartments that were down-regulated in KO vs. WT samples, including synapse organization and transmission, neuron projection, ion transmembrane transport, cation channel complex, synaptic membrane, and somatodendritic compartment (Fig. 7A,B and Supplementary File 2). CX1632-treated KO mice also showed up- regulation of genes associated with glutamatergic and dopaminergic synapses, cAMP, calcium and neuroactive ligand signalling and neuronal system-related functions (Fig. 7C,D).

**Figure 7:**
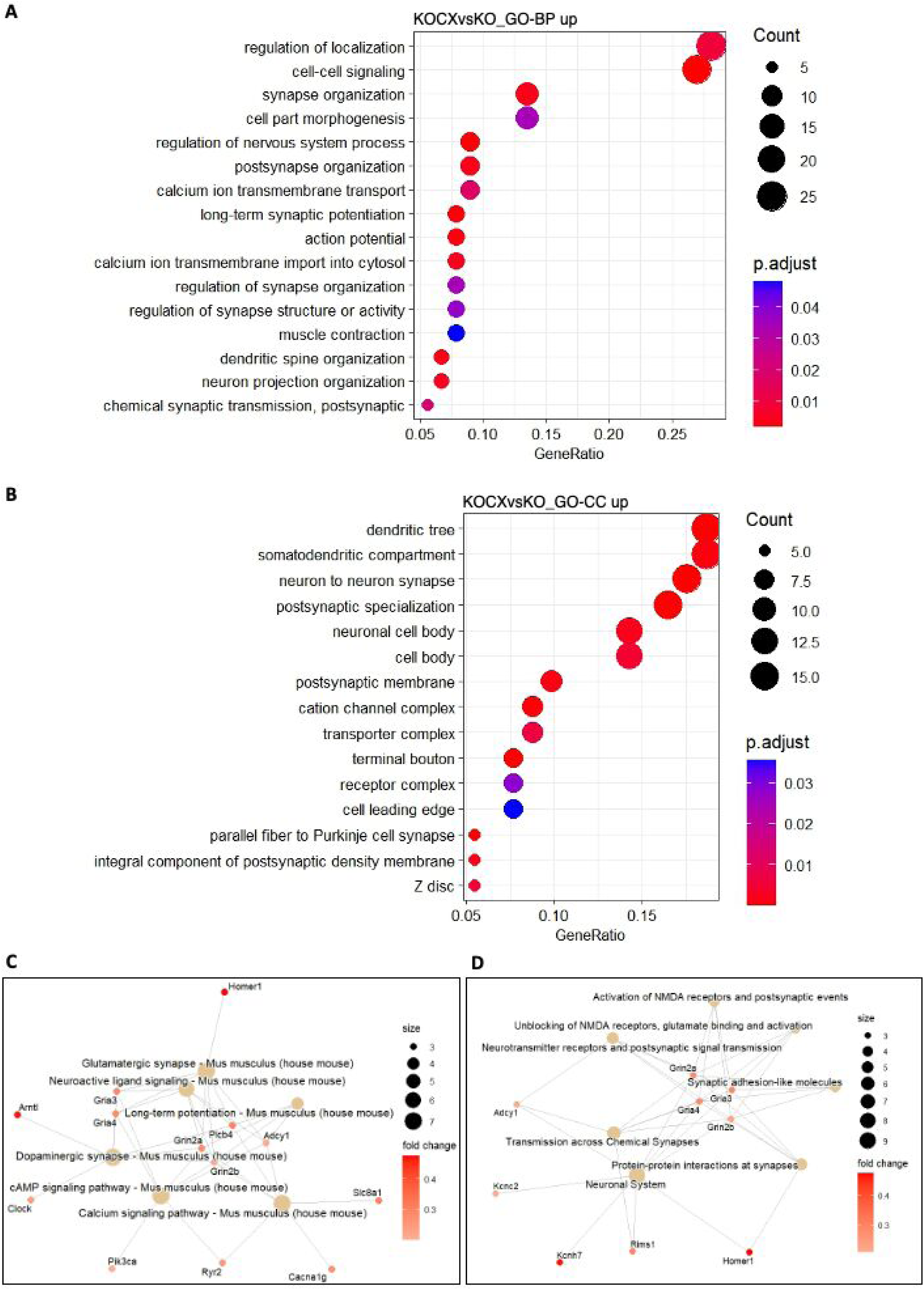
Early CX1632 treatment induces persistent transcriptional remodelling in *Mecp2*-null mice. **(A, B)** Functional enrichment analysis of DEGs identified in CX1632-treated versus vehicle-treated KO mice. Bubble plots show selected Gene Ontology biological process (A) and cellular component (B) terms enriched among upregulated genes. Bubble size indicates the number of contributing DEGs, and color indicates enrichment significance. (**C, D**) Network plots showing upregulated genes contributing to selected KEGG (C) and Reactome (D) pathways. Node size indicates the number of contributing genes, and color indicates gene log₂ fold change.

Taken together, these findings indicate that brief early CX1632 treatment induces persistent transcriptional changes in the *Mecp2*-null prefrontal cortex, that may contribute to the long-lasting behavioral benefits observed after treatment withdrawal.

### CX1632 rescues synaptic organization and restores glutamatergic transmission in *Mecp2-*deficient neurons

Based on the molecular findings, we next tested whether CX1632 could improve synaptogenesis and neuronal activity. WT and KO cortical neurons were treated with CX1632 (1µM) from DIV12 to DIV14, and pre- and postsynaptic puncta density along primary dendrites was assessed by Synapsin 1/2 and Shank2 immunostaining (Fig. 8A). CX1632 rescued the synaptic defects typically observed in KO neurons (Frasca et al, 2024, Pozzer et al, 2025), including reduced puncta density and synaptic markers colocalization, indicating improved synaptic organization (Fig. 8B,C). Consistently, intracellular Ca^2+^ imaging showed a partial but significant rescue of neuronal activity (Fig. 8D,E).

**Figure 8:**
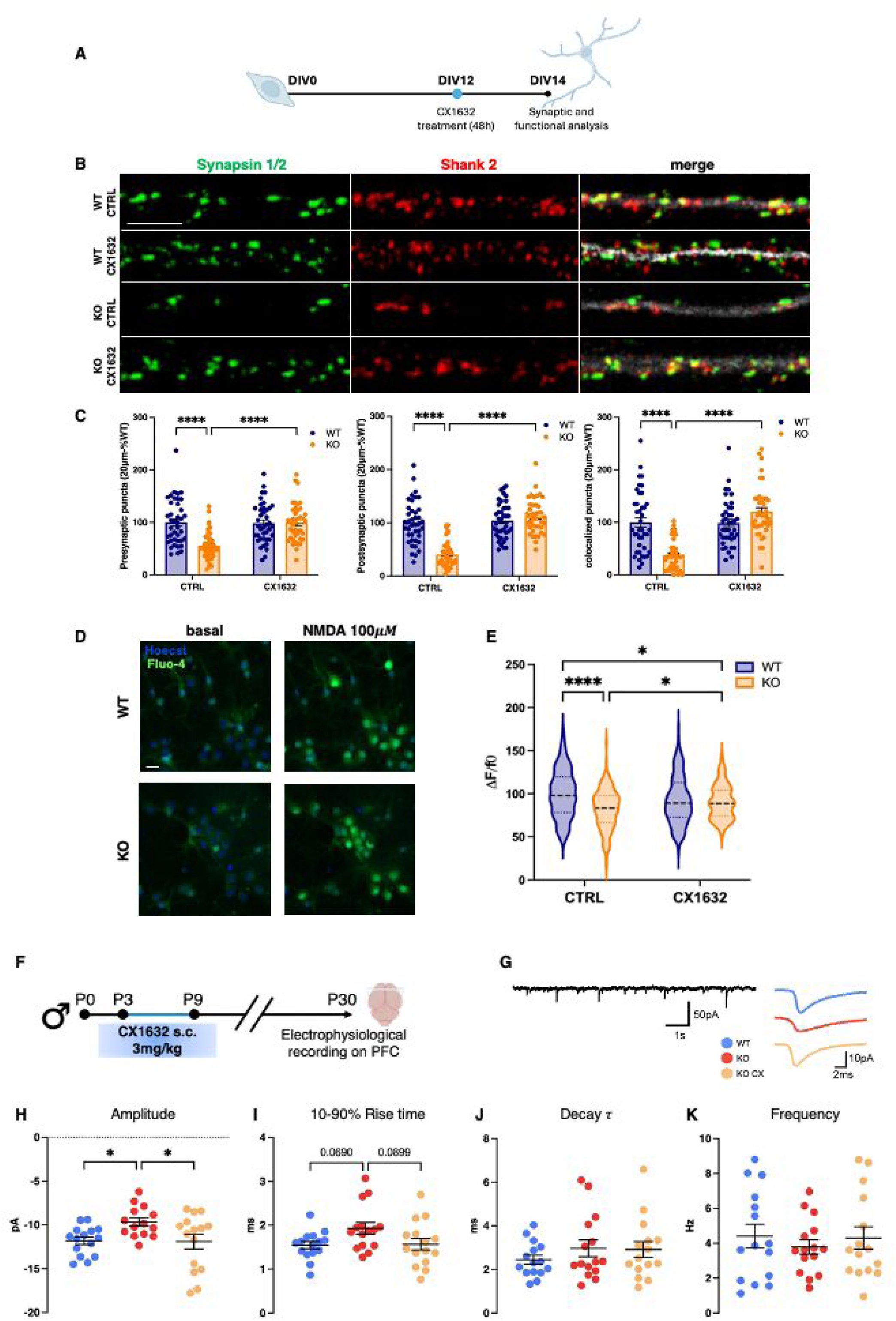
CX1632 rescues synaptic dysfunction in *Mecp2*-deficient neurons. **(A)** Schematic representation of the *in vitro* CX1632 treatment protocol. **(B)** Representative images of Synapsin 1/2 (green), Shank2 (red) and MAP2 (white) immunostaining in DIV14 WT and KO cortical neurons treated with vehicle or 1 μM CX1632 from DIV12 to DIV14. Scale bars, 5 μm. **(C)** Quantification of Synapsin 1/2 and Shank2 puncta density and colocalization. Each dot represents one neuron from three independent neuronal preparations. Statistical significance was assessed by two-way ANOVA followed by Tukey’s multiple-comparisons test. *P < 0.05, **P < 0.01, ***P < 0.001, ****P < 0.0001. Data are presented as mean ± SEM. **(D)** Representative images of Fluo-4-loaded DIV14 WT and KO cortical neurons before and after exposure to 100 μM NMDA. Scale bar, 40 μm. **(E)** Fluo-4 responses, expressed as ΔF/F₀, in vehicle- and CX1632-treated WT and KO neurons. Violin plots show medians and quartiles. Sample sizes were: vehicle-treated WT, n = 193 cells; vehicle-treated KO, n = 148; CX1632-treated WT, n = 159; and CX1632-treated KO, n = 104, from three independent experiments. Statistical significance was assessed by two-way ANOVA followed by Tukey’s multiple-comparisons test. *P < 0.05, **P < 0.01, ***P < 0.001, ****P < 0.0001. **(F)** Schematic representation of the *in vivo* CX1632 treatment and electrophysiological recording protocol. Mice received daily CX1632 or vehicle from P3 to P9, and recordings were performed in prefrontal cortex slices at P30. **(G)** Left, representative voltage clamp recording (excerpted from a 1-min long trace) of sEPSCs in a WT PFC neuron. Right, representative averaged sEPSCs from each experimental group, obtained from 20 individual events. **(H–K)** Quantification of sEPSC amplitude, 10–90% rise time, decay time constant (*τ*) and instantaneous frequency. Vehicle-treated WT, n = 15 cells; vehicle-treated KO, n = 15; and CX1632-treated KO, n = 15, from at least three mice. Statistical significance was assessed by one-way ANOVA followed by Tukey’s multiple-comparisons test. *P < 0.05. Data are presented as mean ± SEM.

To functionally validate the transcriptional data *in vivo*, KO mice were treated with CX1632 from P3 to P9, and patch-clamp recordings were performed at P30 in principal neurons from prefrontal cortex slices obtained from WT, KO and CX1362-treated KO mice (Fig. 8F,G). Spontaneous AMPAR-mediated excitatory postsynaptic currents (EPSCs) were recorded at −70 mV in the presence of the GABA_A_-receptor antagonist gabazine. EPSCs recorded from untreated KO neurons showed significantly reduced amplitude and slower rise time compared with WT neurons, while decay time constant and EPSC frequency were unaffected (Fig. 8H-K). Notably, both EPSC amplitude and rise time were restored in CX1632-treated KO mice 21 days after drug withdrawal (Fig. 8H,I).

Together, these data indicate that CX1632 promotes synaptic maturation and induces a long-lasting functional rescue of glutamatergic transmission in *Mecp2-*deficient neurons.

## Discussion

Experience-dependent neuronal activity shapes brain development by promoting neuronal maturation, synapse stabilization and circuit refinement (Cohen, Greenberg, 2008). Consequently, disrupted activity-dependent signalling may contribute not only to the clinical manifestations of neurodevelopmental disorders but also to the establishment of abnormal circuit trajectories (Ebert, Greenberg 2013). This concept is particularly relevant to RTT, in which impaired neuronal maturation, glutamatergic transmission and network activity emerge before overt neurological symptoms (Kishi, Macklis, 2004, Bedogni et al, 2016, Scaramuzza et al, 2021, Illescas et al, 2024, Osaki et al, 2026). Consistently, environmental, circuit-directed and pharmacological interventions that modify neuronal activity or plasticity have produced beneficial effects in *Mecp2-*deficient models, indicating that RTT circuits retain a therapeutically exploitable capacity for functional remodeling (Kron et al, 2009, Degano et al, 2014, Scaramuzza et al, 2021, Achilly et al, 2021, Lu et al, 2016, Hao et al, 2015, Patrizi et al, 2016).

Here, we show that positive allosteric modulation of AMPA receptors with CX1632 produces sustained therapeutic effects in *Mecp2*-deficient mice and that efficacy is strongly influenced by developmental stage and administration schedule. A brief treatment from P3 to P9 delayed disease progression, prolonged survival and improved selected motor and cognitive outcomes in *Mecp2*-null males, with persistent benefits also observed in heterozygous females. Repeated intermittent administration initiated during the same early developmental period further strengthened or prolonged selected effects, whereas a brief treatment at a later juvenile stage showed markedly lower efficacy, particularly in already symptomatic null males. Although our previous study with CX546 provided initial evidence that early AMPAR potentiation can produce long-lasting effects^21^, to our knowledge, no previous pharmacological treatment in an RTT model has produced such broad benefits persisting for several weeks to months after only seven days of administration. The present findings therefore reveal an unprecedented breadth and duration of efficacy using a pharmacologically more advanced AMPAR modulator (Kadriu et al, 2021, Bretin et al, 2017, Freudenberg et al, 2025, Giralt et al, 2017).

CX1632, also known as S47445 or Tulrampator, is a potent AMPAR-PAM with procognitive, neurotrophic and synaptic plasticity-enhancing properties in preclinical models (Bretin et al, 2017, Freudenberg et al, 2025, Giralt et al, 2017). Previous phase II evaluation showed tolerability during prolonged administration in humans, although no significant efficacy was detected in patients with mild-to-moderate Alzheimer’s disease and depressive symptoms, supporting its translational relevance for RTT (Kadriu et al, 2021). Unlike direct receptor agonists, AMPAR-PAMs enhance responses to endogenous glutamate and should therefore preserve the spatial and temporal organization of physiological synaptic signalling. We found that CX1632 increased GluA1 Ser845 and Akt phosphorylation in cortical neurons, supporting engagement of AMPAR-associated signalling. Moreover, comparable cortical GluA1 abundance and phosphorylation in WT and *Mecp2*-null mice indicate that the *Mecp2*-deficient brain retains molecular substrates potentially responsive to AMPAR modulation, even after symptoms have begun to emerge.

The use of both *Mecp2*-null males and heterozygous females was important because the two models provide complementary information. Heterozygous females more closely reproduce the genetic mosaicism of typical RTT and the interactions between MeCP2-positive and MeCP2-negative cells. However, the milder, more heterogeneous, and more slowly progressive phenotype of female mice, compared with that observed in many patients and in KO mice, reduces the sensitivity of this model to detect treatment effects. Conversely, null males do not reproduce the mosaic genetic condition of typical RTT but develop a severe and temporally synchronized phenotype that facilitates the analysis of survival and disease progression. Evaluating both models therefore allowed us to address both genetic relevance and phenotypic severity, although direct quantitative comparisons between sexes are not possible.

The most striking result was the persistent efficacy of CX1632 when administered during the first postnatal week. P3–P9 encompasses a period of intense synaptic and circuit maturation in mice (Hooks, Chen, 2007) and broadly overlaps, although not through a direct age equivalence, with late prenatal-to-early neonatal brain development in humans (Semple et al, 2013). Treatment during this window produced benefits that persisted for weeks to months after drug withdrawal, suggesting that transient enhancement of excitatory transmission redirected the subsequent developmental trajectory of *Mecp2*-deficient circuits. This finding is particularly relevant as expanding genomic newborn screening initiatives may increasingly identify children carrying pathogenic *MECP2* variants before overt regression (Singh, Santosh, 2024), potentially creating opportunities for presymptomatic intervention.

The stronger and earlier improvement produced by repeated intermittent treatment further indicates that efficacy depends not only on treatment onset but also on exposure pattern. Intermittent administration may repeatedly reinforce activity-dependent maturation while avoiding uninterrupted pharmacological stimulation. However, the current experiments do not establish the optimal duration of treatment cycles or off-treatment intervals. Whether shorter intervals, different doses or alternative schedules could further increase efficacy without compromising tolerability will require systematic pharmacokinetic and pharmacodynamic optimization.

The limited effect of a brief P28–P34 treatment in null males supports the existence of a developmentally restricted component of therapeutic responsiveness. At this stage, *Mecp2*-null males already display mild overt symptoms and have experienced several weeks of abnormal neuronal development. Consequently, AMPAR potentiation may be less able to redirect circuit organization once dysfunctional connectivity has become progressively established. Nevertheless, the partial efficacy of prolonged intermittent treatment initiated at P20 indicates that the juvenile *Mecp2*-null brain is not completely refractory to modulation. Moreover, the response to late treatment observed in heterozygous females suggests that chronological age alone is unlikely to determine efficacy. Rather, therapeutic responsiveness may depend on the interaction between residual developmental plasticity and disease burden, including the extent to which altered circuits have stabilized. Because null males and heterozygous females differ in sex, *Mecp2* dosage, cellular mosaicism and disease kinetics, the present experiments cannot formally disentangle the contribution of developmental age from that of pathological progression.

These considerations are relevant to clinical translation but should not be interpreted as defining a direct age limit for treatment in patients. Murine developmental stages cannot be converted into specific human ages using a single linear relationship (Sempl et al, 2013, Dutta, Sengupta, 2016). Our data instead suggest that disease stage, residual circuit plasticity and treatment schedule should be considered when designing trials of activity-dependent therapies. They also raise the possibility that interventions targeting neuronal activity may be most effective before circuit dysfunction becomes extensively consolidated, while still retaining some benefit at later stages in more slowly progressing or less severely affected conditions.

The persistence of behavioral effects long after drug withdrawal suggested that CX1632 induced changes extending beyond its acute pharmacological activity. To investigate this possibility, we analyzed the transcriptomic profile of the prefrontal cortex at P30, 21 days after completion of neonatal treatment. At this mildly symptomatic stage, *Mecp2*-null mice displayed relatively modest transcriptional dysregulation; nevertheless down-regulated genes converged on functional related processes involving neuronal development, synaptic organization and signaling, ion transport and LTP potentiation. In CX1632-treated KO mice, enriched processes were predominantly up-regulated and involved many of the same functional domains impaired by *Mecp2* deficiency, including synapse organization, glutamatergic signaling, calcium and ion transport, action potential regulation and LTP-associated processes. Thus, although the transcriptomic analysis was based on a limited number of treated KO samples following conservative exclusion of a litter showing distinct clustering, the direction and functional coherence of the response suggest that early CX1632 administration persistently opposes disease-associated pathway alterations rather than merely producing an acute transcriptional effect.

The biological relevance of this transcriptional signature is further supported by its convergence with independent cellular and functional findings. In *Mecp2*-deficient cortical cultures, CX1632 rescued pre- and postsynaptic puncta density and colocalization and partially restored neuronal calcium responses, consistent with enhanced synaptic maturation and activity. Most importantly, electrophysiological recordings demonstrated that neonatal treatment produced a persistent rescue of AMPAR-mediated transmission in prefrontal cortical neurons, normalizing the reduced EPSC amplitude and slower rise time observed in untreated *Mecp2*-null mice 21 days after drug withdrawal. These results provide functional validation of the pathways identified by RNA sequencing and indicate that the molecular effects of CX1632 translate into durable improvements in synaptic organization and cortical neuronal function.

In conclusion, our findings identify developmental timing as a major determinant of the therapeutic response to AMPAR potentiation in RTT models. Brief CX1632 administration during an early period of circuit maturation produced broad and persistent benefits, which could be strengthened by intermittent re-exposure, whereas later treatment was less effective and more dependent on disease stage and treatment schedule. By linking sustained behavioral efficacy to persistent transcriptional, synaptic and electrophysiological changes, this study supports the concept that transient enhancement of activity-dependent signaling can modify the developmental trajectory of *Mecp2*-deficient circuits. These findings provide a rationale for evaluating AMPAR positive allosteric modulators as developmentally informed therapeutic strategies, particularly in the context of increasingly early genetic diagnosis of *MECP2*-related disorders.

## Materials and methods

### Animals

The *Mecp2* mouse strain was originally purchased from Jackson Laboratories (B6.129P2(C)-Mecp2^tm1.1Bird^/J) and backcrossed onto a CD1 genetic background (Cobolli Gigli et al, 2016). Mice were generated by crossing *Mecp2^+/−^* females with wild-type male mice purchased from Charles

River Laboratories. Genotyping was performed as previously described^21^. For primary neuronal cultures, the day of vaginal plug was considered E0.5.

All procedures complied with Directive (2010/63/UE) and were approved by the Italian Ministry of Health (authorizations n° 1172/2020-PR and n° 33/2024-PR) and the San Raffaele Scientific Institutional Animal Care and Use Committee.

### Primary cortical neurons

Primary cortical neurons were prepared from E15.5 mouse embryos. Cerebral cortices were rapidly dissected and kept in cold HBSS (Life Technologies, #4175-095) until dissociation. Tissues were incubated with 0.25% trypsin/EDTA (GIBCO, Thermo Fisher Scientific) for 10 min at 37°C, and digestion was blocked with DMEM (Life Technologies, #41966-029) containing 10% FBS (Thermo Fisher Scientific, #10500064). Cortices were then mechanically dissociated in DMEM containing 10% FBS, 1% L-glutamine (Sigma-Merck, #G7513), and 1% penicillin/streptomycin (P/S; Sigma-Merck, #P0781).

Depending on the experiment, neurons were seeded on poly-D-lysine-coated plates (0.1 or 1 mg/mL; Sigma-Merck, #P7886) or glass coverslips (Neuvitro, #GG-12-PDL) in neuronal medium consisting of Neurobasal Plus medium (Thermo Fisher Scientific, #A3582901), 1% P/S, and 2% B27 Plus 50X (Thermo Fisher Scientific, #A3582801).

CX1632 was added directly to the culture medium at 12 days *in vitro* (DIV12) or DIV14 at 0.75, 1.5, or 5 μM; control cultures received corresponding volumes of neuronal culture medium.

### Western blotting of neuronal extracts

Total proteins were extracted from DIV14 cortical neurons using ice-cold RIPA buffer (100 mM Tris-HCl pH 7.5, 300 mM NaCl, 10 mM EDTA, 2% NP-40, 0.2% SDS, 1% sodium deoxycholate, 1X Protease Inhibitor Cocktail (Thermo Fisher Scientific, #78444), and 1X PhosSTOP (Sigma-Merck, #4906845001). Protein concentration was determined using the bicinchoninic acid (BCA) assay kit (Thermo Fisher Scientific, #23228), according to the manufacturer’s instructions.

Protein lysates (20 μg) were separated on TGX Stain-Free gels (Criterion, 12/18 wells, 4–15%; Bio-Rad, #5678084) and transferred onto nitrocellulose membranes (Trans-Blot Turbo Nitrocellulose Midi; Bio-Rad, #1704159). After transfer, membrane images were acquired with the Bio-Rad ChemiDoc system and used for data normalization. Membranes were blocked 1 h at room temperature and incubated overnight (4°C) with the following primary antibodies diluted 1:1000 in blocking solution: anti-AKT (Cell Signaling, #4685;), anti-phospho-AKT Ser473 (Cell Signaling, #4060), anti-GluA1 (Cell Signaling, #13185), and anti-phospho-GluA1 Ser845 (Abcam, #76321).

After three washes in 1X TBS-T, membranes were incubated with the appropriate HRP-conjugated secondary antibodies: peroxidase-conjugated AffiniPure goat anti-rabbit IgG (H+L) (#111-035-144) or goat anti-mouse IgG (H+L) (#115-035-003; Jackson ImmunoResearch). Immunocomplexes were detected using ECL substrates from Cyanagen (WESTAR SUN, #XLS0630250) or Bio-Rad (Clarity Western ECL Substrate, #1705061) and imaged with the ChemiDoc System. Band quantification was performed using Image Lab 5.2.1 software (Bio-Rad).

### TIF preparations and analysis

Mutant and WT littermates were sacrificed by cervical dislocation and brains were rapidly removed. Cortices were dissected, snap-frozen on dry-ice and stored at -80°C until use. Triton-insoluble fractions (TIF), enriched in pre- and post-synaptic markers, and cytosolic fractions were obtained as in Gardoni et al., 1998. Briefly, frozen cortices were homogenized in ice-cold lysis buffer (320 mM sucrose, 1 mM NaHCO_3_, 1 mM HEPES pH 7.4, 1 mM MgCl_2_, protease inhibitor cocktail (Thermo Fisher scientific #78444) and PhosSTOP 1X and centrifuged to isolate nuclei. Supernatants, that contain the cytosolic fraction, were collected and centrifuged again 13000g at 4oC for 15 minutes. Pellets were resuspended in KCl triton buffer (containing KCl 150, Triton 0.5%) and then centrifuged at 100000g (Ultracentrifuge: Optima TLX 1200000rpm, Beckman Coulter; rotor: Beckman TLA-55 07U) at 4°C for 1 hour. Then, the obtained pellet was homogenized, using a potter in glass, in T-PER (Thermo Scientific #78510). Protein quantification and western blotting were performed as described above.

### Analysis of synaptic puncta

For immunofluorescence, DIV14 neurons grown on glass coverslips were fixed in 4% PFA/10% sucrose in PBS, permeabilized with 0.2% Triton X-100 in PBS, blocked with 4% BSA in PBS, and incubated with primary antibodies diluted in 0.2% BSA in PBS: anti-MAP2 1:1000 (clone D5G1; Cell Signaling, #8707), anti-Synapsin1/2 1:500 (Synaptic Systems, #106006), and anti-Shank2 1:300 (Synaptic Systems, #162211). Cells were then incubated with Alexa Fluor-conjugated secondary antibodies, counterstained with DAPI (Thermo Fisher Scientific, #D1306), and mounted with Fluoromount (Thermo Fisher Scientific, #00-4958-02).

For synaptic puncta density and colocalization analyses, Z-stacks were acquired with a Leica Stellaris 8 DLS microscope equipped with an HC PL APO CS2 63×/NA 1.4 oil-immersion objective and three HyD S detectors. Images were acquired at 1024 × 1024 pixels, 16-bit depth, over 127.28 × 127.28 μm² fields, with a 0.3 μm step size, using a 405 nm diode laser and a pulsed tunable white-light laser. Acquisition parameters were kept constant within each experiment.

Puncta density was quantified in ImageJ by counting synaptic puncta ≥0.16 μm² within manually selected ROIs along primary dendrites (20 μm, three branches/neuron). Pre- and post-synaptic marker colocalization was assessed with the Colocalization Highlighter plugin on maximum-intensity projection binary masks. Colocalized puncta were quantified in manually selected ROIs, and only puncta ≥0.1 μm² were counted (Frasca et al, 2020, Albizzati et al, 2024).

### Calcium imaging

DIV14 primary cortical neurons were loaded with 2 μM Fluo-4 (Invitrogen, #F14201) in KRH buffer (125 mM NaCl, 5 mM KCl, 1.2 mM MgSO₄, 1.2 mM KH₂PO₄, 25 mM HEPES, 6 mM glucose, 2 mM CaCl₂, pH 7.4) for 30 min at 37°C and washed once with the same buffer.

NMDA stimulation was performed using the liquid-handling system of the ArrayScan XTI HCA Reader (Thermo Fisher Scientific) by adding 100 μM NMDA at 50 μL/s during image acquisition. Images were acquired with a high-resolution Photometrics camera through a 20× objective (Zeiss Plan-NEOFLUAR, 0.4 NA). Fluo-4 and Hoechst signals were recorded for 40 frames at 1 Hz, with exposure times of 40 and 25 ms, respectively; at least nine baseline frames were acquired before stimulation.

Analysis was performed with HCS Studio software using the SpotDetector bioapplication (Thermo Fisher Scientific). Hoechst-positive nuclei were identified, and Fluo-4 mean intensity was measured in the soma after background subtraction. Only cells with neuronal morphology were included. Calcium responses were expressed as ΔF/F₀.

### Pharmacological treatments

Ampakine CX1632 (3 mg/kg; MedChemExpress, #HY-109046) was administered daily by subcutaneous injection from P3 to P9 (50 μL) or by intraperitoneal injection in adult mice. CX1632 was dissolved in 10% DMSO, 5% Tween-80, 40% PEG400, and 45% saline. Control mice received vehicle only.

### RNA purification

P30 mutant mice and WT mice were euthanized by cervical dislocation, and brains were rapidly removed. Prefrontal cortices were dissected, snap-frozen on dry ice, and stored at −80°C until use. Total RNA was extracted with PureZol (Bio-Rad, #7326890), followed by DNase I treatment to remove genomic DNA (Sigma, #AMPD1). RNA integrity was assessed using a Fragment Analyzer (Agilent Technologies), and RNA concentration was measured with a Qubit Fluorometer (Thermo Fisher Scientific). All 24 samples, six per experimental group, met the requirements for RNA sequencing (RNA integrity number, RIN >7) and were further processed.

### RNA sequencing, library preparation and bioinformatics data analysis

PolyA+ RNA-seq libraries were prepared from six animals per experimental group using the NEBNext Ultra II RNA Library Preparation Kit (New England Biolabs), according to the manufacturer’s instructions. Libraries were sequenced by GENEWIZ/Azenta Life Sciences on an Illumina NovaSeq X Plus platform using a 2 × 150-cycle run, generating at least 30 million fragments per sample.

Raw reads were aligned, counted, and quality-controlled using the nf-core/rnaseq pipeline (v3.18.0) (De Keyzer et al, 2026, Ewels et al, 2020) running in Nextflow (v24.10.5) (Di Tommaso et al, 2017), with default parameters. The GRCm39 mouse genome and GENCODE M38 annotation were used as references. Salmon gene-level counts without offset or bias correction were used for differential expression analysis with DESeq2 v1.48.1 in R (v4.5.1) (Love et al, 2014). Outlier samples were removed before counts normalization and only expressed genes, defined as genes with a total read count ≥10 in more than 3 samples, were maintained for statistical significance. Genes with |log₂ fold change| >0.2 and Benjamini–Hochberg-adjusted p value <0.05 were considered differentially expressed genes (DEGs).

Functional enrichment over-representation analysis of Gene Ontology terms and KEGG/Reactome pathways was performed using clusterProfiler R package (v4.6.2) (Yu et al, 2012). Enrichments with Benjamini–Hochberg-adjusted false discovery rate (FDR) <0.05 were considered significant.

Raw sequence data and gene counts are available in the Gene Expression Omnibus repository under accession number GSE338385.

### Patch clamp recordings

P30 mice were anaesthetized with ketamine/xylazine (100 and 10 mg/kg, i.p.) and transcardially perfused with ice-cold artificial cerebrospinal fluid (ACSF) containing 125 mM NaCl, 3.5 mM KCl, 1.25 mM NaH2PO4, 2 mM CaCl2, 25 mM NaHCO3, 1 mM MgCl2, and 11 mM D-glucose, saturated with 95% O2/5% CO2, pH 7.3. After decapitation, brains were removed and 300-μm coronal slices containing the prefrontal cortex (PFC) were cut in ice cold ACSF using a VT1000S vibratome (Leica Microsystems). Slices were in a recording chamber mounted on an upright BX51WI microscope (Olympus) equipped with differential interference contrast optics and perfused with ACSF containing 10 µM SR95531 (gabazine, Abcam, #ab120042) at 2–3 mL/min and 32°C. Whole-cell patch-clamp recordings were performed in PFC pyramidal cells using micropipettes filled with 10 mM NaCl, 124 mM KH2PO4, 10 mM HEPES, 0.5 mM EGTA, 2 mM MgCl2, 2 mM Na2-ATP and 0.02 mM Na-GTP (pH 7.2, adjusted with KOH; tip resistance: 4–6 MΩ). AMPA-receptor mediated spontaneous excitatory postsynaptic currents (sEPSCs) were recorded in voltage clamp at a holding potential of −70 mV for 1 min. Signals were acquired with a MultiClamp 700B amplifier and Digidata 1440A digitizer using pClamp10 software (Molecular Devices), sampled at 10 kHz and low-pass filtered at 2 kHz. sEPSC amplitude, 10-90% rise time, decay time constant, and instantaneous frequency were analysed with Clampfit and GraphPad Prism; median values from each trace were used for statistical comparisons.

### Behavioural assessment

Mice were maintained under an inverted 12-h light/dark cycle at 22–24°C. Behavioural testing was performed during the dark phase by investigators blinded to genotype and treatment. Sample sizes are reported in the corresponding figure legends.

### Phenotypic scoring

Body weight and phenotypic severity were assessed twice weekly from P20 in males and weekly from P60 in females. Phenotypic scoring was performed as previously described^25,35^ and included general condition, mobility, hindlimb clasping, tremor and gait. Mice scoring 2 for general condition or tremor, or showing rapid weight loss, were euthanized for ethical reasons. The day of euthanasia was considered the survival endpoint.

### Open-field test

Exploratory behaviour and locomotor activity were assessed in a 40 × 40 cm open-field arena. Each mouse was placed individually in the center of the arena and allowed to explore freely for 10 min. Total distance travelled was measured using EthoVision XT 14 software (Noldus) as an index of locomotor activity.

### Rotarod test

Motor coordination and motor learning were assessed using an accelerating rotarod apparatus (Ugo Basile, Stoelting Co.). The test was performed over three consecutive days, comprising two training days and one test day. Each daily session consisted of three 5-min trials. Rotation speed increased progressively from 4 to 40 rpm. Each trial ended when the mouse fell or after 5 min, and latency to fall was recorded automatically.

### Pole test

Motor coordination and balance were assessed using the pole test. Mice underwent three training trials before testing. Each mouse was placed near the top of a vertical pole with its head facing upward and allowed to turn downward and descend towards its home cage. Total time required to orient and reach the base of the pole was recorded over three trials, with a maximum duration of 60 s per trial.

### Beam-walking test

Balance and fine motor coordination, particularly hindlimb function, were assessed using the beam-walking test. Mice were trained to traverse a narrow beam connecting a starting platform to their home cage. The beam was 1–3 cm wide and elevated above the floor. During testing, traversal time and the number of foot slips were recorded.

### Spontaneous alternation test

Spontaneous alternation was assessed in a four-arm maze as a measure of exploratory behaviour and spatial working memory. Each mouse was placed in the center of the maze and allowed to explore freely for 10 min. An arm entry was scored when all four limbs entered the arm. The sequence and total number of arm entries were recorded. Alternation percentage was calculated as:

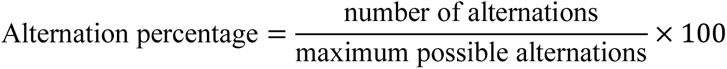

### Novel object recognition test

The novel object recognition test was used to assess recognition memory. During the training session, each mouse was allowed to explore two identical objects for 10 min. After a 30-min retention interval, one familiar object was replaced with a novel object, and mice were allowed to explore both objects. Object exploration was scored by an investigator blinded to genotype and treatment. The discrimination index was calculated as:

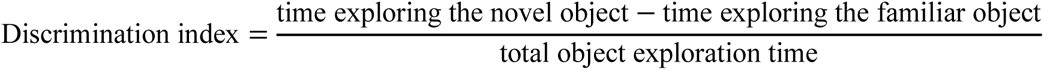

### Marble-burying test

Environment-directed exploratory behaviour was assessed using the marble-burying test. Each mouse was placed individually in a cage containing 5 cm of standard bedding. Twelve glass marbles, 19 mm in diameter, were evenly arranged on the bedding in four rows of three. Mice were allowed to explore for 30 min, after which marbles buried to at least two-thirds of their depth were counted.

### Novelty-exposure test

The test consisted of two sessions. During the first session, mice were allowed to explore an empty arena freely for 10 min. Immediately after, an object was placed in the center of the same arena, and mice were allowed to explore it for 10 min. Object-directed behaviour was assessed by measuring exploration time and the number of discrete exploratory bouts. Locomotor activity during habituation was evaluated as the total distance travelled.

Effect sizes for each behavioral test are reported in Table 1.

**Table 1.** The effect sizes Hedges’ g for behavioral tests performed on male and female mice.

| Male mice | WT vs KO | KO vs KO CX1632 | KO over time | KO CX1632 over time |
| --- | --- | --- | --- | --- |
| Phenotypic score P80 (Fig. 2C) | **6,864065** | **1,5** |  |  |
| Phenotypic score P90 (Fig. 2C) | **12,222326** | **2,090438** |  |  |
| Rotarod test P45 (Fig. 2D) | **4,59491** | **0,995594** |  |  |
| Motor learning P45 (Day1 vs 2, Fig. 2D) |  |  | 0,637861 | **0,871831** |
| Motor learning P45 (Day1 vs 3, Fig. 2D) |  |  | 0,552844 | **0,819636** |
| Pole test P65 (Fig. 2E) | **2,332037** | **0,985817** |  |  |
| Improvement Pole test (Fig. 2E) | **1,26316** | **0,956007** |  |  |
| Novel Object Recognition P45 (Fig. 2G) | **1,75371** | **1,02785** |  |  |
| Motor learning P45 (Day1 vs 2, Fig. 3I) |  |  | 0,75472 | **0,992946** |
| Motor learning P45 (Day1 vs 3, Fig. 3I) |  |  | **1,044743** | **1,212553** |
| Beam walking test P60 (Fig. 3J) | **1,39180** | 0,479522 |  |  |
| Pole test P41 (Fig. 3K) | 0,243094 | 0,300254 |  |  |
| X maze P45 (Fig. 3L) | **1,43414** | 0,672431 |  |  |
| Marble Burying test P62 (Fig. 3M) | **2,12607** | **1,25419** |  |  |
| Phenotypic score P60 (Fig. 5C) | **5,176592** | **2,162922** |  |  |
| Phenotypic score P70 (Fig. 5C) | **13,609555** | **2,00069** |  |  |
| Phenotypic score P80 (Fig. 5C) | **11,110056** | **3,386669** |  |  |
| Rotarod test P45 (Fig. 5D) | **4,05592** | **1,25774** |  |  |
| Motor learning P45 (Day1 vs 2, Fig. 5D) |  |  | 0,339347 | **1,035685** |
| Motor learning P45 (Day1 vs 3, Fig. 5D) |  |  | **0,179693** | **0,938488** |
| Rotarod test P65 (Fig. 5D) | **12,05545** | **1,39926** |  |  |
| Pole test P65 (Fig. 5E) | **3,23575** | **1,62046** |  |  |
| Novel Object Recognition P65 (Fig. 5G) | 0,24183 | **1,038341** |  |  |
| Female mice | WT vs HET | HET vs HET CX1632 | HET over time | HET CX1632 over time |
| Novel Object Recognition P45 (Fig. 2J) | 0,475555 | **0,895355** |  |  |
| Rotarod test P100 (Fig. 2K) | **5,498697** | **2,0458006** |  |  |
| Motor learning (Day1 vs 2, Fig. 2K) |  |  | **0,816801** | **1,151754** |
| Motor learning (Day1 vs 3, Fig. 2K) |  |  | **1,008852** | **1,316645** |
| Foot slips errors P100 (Fig. 2L) | **1,320516** | **1,055262** |  |  |
| Rotarod test P100 (Fig. 4C) | **1,746251** | 0,757028 |  |  |
| Motor learning (Day1 vs 2, Fig. 4C) |  |  | 0,776129 | **1,280755** |
| Motor learning (Day1 vs 3, Fig. 4C) |  |  | **0,894323** | **0,92633** |
| Foot slips errors P100 (Fig. 4D) | **1,086413** | 0,58924 |  |  |
| Novelty time P115 (Fig. 4E) | **1,188709** | 0,610614 |  |  |
| Open Field P115 (Fig. 4F) | **1,258325** | 0,072689 |  |  |

Only behavioral tests showing improvement/rescue are included in the effect size analysis reported in the table. 0 < enrichment score (ES) < 0.20 Ignored; 0.20 < ES < 0.50 small; 0.50 < ES < 0.80 moderate; 0.80 < ES < 1.30 large; 1.30 > ES Very Large. Large and very large are indicated in bold. HET = heterozygous; KO = knockout; WT = wild type, P = postnatal day.

### Statistical analysis

Statistical analysis and data plotting were performed using GraphPad Prism 10. Pairwise comparisons were analysed using t-test or Mann-Whitney U test, as appropriate. Comparisons involving three or four experimental groups were analyzed by two-way or one way-ANOVA respectively, to assess the effects of genotype, ampakine treatment and their interaction. When there was a significant effect of treatment or genotype, or a significant interaction between the variables, Tukey’s post-hoc test was applied. Statistical significance was set at p-value < 0.05. Outliers were identified using the ROUT test and excluded from the analysis. Effect size was calculated using Hedges’ g. Culture wells and mice were randomly assigned to treatments in *in vitro* and *in vivo* experiments, respectively. Investigators were blinded to treatment and genotype during data acquisition and analysis.

## Data availability

RNA-sequencing datasets are available at the Gene Expression Omnibus (GEO) database; accession number GSE338385.

## Author contributions

**Giuseppina De Rocco:** Conceptualization; Investigation; Data curation; Formal analysis; Visualization, Writing–original draft. **Andrea de Donato**: Investigation; Formal analysis**. Marzia Indrigo:** Investigation. **Virginia Varotto:** Investigation. **Martina Geusa**: Investigation. **Stefano Taverna:** Investigation; Data curation; Formal analysis; Writing–original draft. **Ingrid Cifola**: Data curation, Software; Investigation; Bioinformatic analysis, Writing–original draft**. Eva Maria Pinatel**: Data curation, Investigation; Bioinformatic analysis, Writing–original draft. **Angelisa Frasca**: Conceptualization, Formal analysis. **Nicoletta Landsberger**: Conceptualization; Visualization; Supervision; Funding acquisition; Writing–original draft.

## Supplementary material

Supplementary material is available at EMBO Molecular Medicine online.

## Acknowledgements

We are grateful to Prof. Fabrizio Gardoni and Dr. Maria Italia (Università degli Studi di Milano) for sharing protocols, suggestions and reagents and to Dr. Patrizia D’Adamo and the Mouse Behaviour facility (IRCCS, San Raffaele Scientific Institute) for assistance in behavioural tests. We are grateful to Dr. Francesco Bedogni for valuable discussions during the development of this project and to Dr. Tilo Schorn for technical assistance with confocal imaging. We thank Dr. Maria Balbontin Arenas for her availability and support with bioinformatic analyses. We acknowledge: ALEMBIC (Advanced Light and Electron Microscopy BioImaging Center, IRCCS San Raffaele) and specifically Desiree Zambroni for assistance in calcium imaging experiments. We thank all the master’s and MD students who contributed to this project over the years for their valued support. Finally, we gratefully acknowledge the longstanding financial and emotional support provided to our laboratory by ProRETT Ricerca, an Italian parents’ association.

## Expanded View Figure Legends

**Figure EV1: Longitudinal analysis of individual phenotypic severity components and locomotor activity. (A-U)** Line graphs show longitudinal scores for individual phenotypic parameters, including general condition, mobility, hindlimb clasping, tremor and gait. (A-E) Male mice treated daily with CX1632 or vehicle from P3 to P9. (F) Distance travelled by female mice during the open-field test at P90. (G-K) Male mice treated daily from P28 to P34. (L-P) Male mice receiving intermittent treatment from P20 to P68. (Q-U) Male mice receiving repeated seven-day treatment cycles initiated at P3 and separated by two-week off-treatment intervals.

**Figure EV2: RNA-seq quality control and functional alterations in *Mecp2*-null prefrontal cortex. (A)** Principal component analysis of all sequenced samples: six vehicle-treated WT, six vehicle-treated KO, six CX1632-treated WT and three CX1632-treated KO mice. **(B)** Bubble plot showing selected Gene Ontology cellular component terms enriched among downregulated genes. Bubble size indicates the number of contributing DEGs, and color indicates enrichment significance. **(C)** Network plot showing downregulated genes contributing to selected Reactome pathways. Node size indicates the number of contributing genes, and colour indicates gene log₂ fold change

